# An adjuvanted SARS-CoV-2 RBD nanoparticle elicits neutralizing antibodies and fully protective immunity in aged mice

**DOI:** 10.1101/2021.09.09.459664

**Authors:** Francesco Borriello, Etsuro Nanishi, Hyuk-Soo Seo, Timothy R. O’Meara, Marisa E. McGrath, Yoshine Saito, Robert E. Haupt, Jing Chen, Joann Diray-Arce, Kijun Song, Andrew Z Xu, Timothy M. Caradonna, Jared Feldman, Blake M. Hauser, Aaron G. Schmidt, Lindsey R. Baden, Robert K. Ernst, Carly Dillen, Stuart M. Weston, Robert M. Johnson, Holly L. Hammond, Jingyou Yu, Aiquan Chang, Luuk Hilgers, Peter Paul Platenburg, Sirano Dhe-Paganon, Dan H. Barouch, Al Ozonoff, Ivan Zanoni, Matthew B. Frieman, David J. Dowling, Ofer Levy

**Affiliations:** Division of Immunology, Boston Children’s Hospital, Boston, MA, USA; *Precision Vaccines Program*, Division of Infectious Diseases, Boston Children’s Hospital, Boston, MA, USA; Department of Pediatrics, Harvard Medical School, Boston, MA, USA; Department of Cancer Biology, Dana-Farber Cancer Institute, Boston, MA, USA; Department of Biological Chemistry and Molecular Pharmacology, Harvard Medical School, Boston, MA, USA; Department of Microbiology and Immunology, University of Maryland School of Medicine, Baltimore, MD, USA; Research Computing Group, Boston Children’s Hospital, Boston, MA, USA; Ragon Institute of MGH, MIT, and Harvard, Cambridge, MA, USA; Department of Microbiology, Harvard Medical School, Boston, MA, USA; Department of Medicine, Brigham and Women’s Hospital, Boston, MA, USA; Department of Microbial Pathogenesis, University of Maryland School of Dentistry, Baltimore, MD, USA; Center for Virology and Vaccine Research, Beth Israel Deaconess Medical Center, Harvard Medical School, Boston, MA, USA; LiteVax B.V., Oss, The Netherlands; Broad Institute of MIT & Harvard, Cambridge, MA, USA

**Keywords:** SARS-CoV-2, COVID-19, vaccine, nanoparticle, RBD, adjuvant, innate immunity, antibody

## Abstract

Development of affordable and effective vaccines that can also protect vulnerable populations such as the elderly from COVID-19-related morbidity and mortality is a public health priority. Here we took a systematic and iterative approach by testing several SARS-CoV-2 protein antigens and adjuvants to identify a combination that elicits neutralizing antibodies and protection in young and aged mice. In particular, SARS-CoV-2 receptor-binding domain (RBD) displayed as a protein nanoparticle (RBD-NP) was a highly effective antigen, and when formulated with an oil-in-water emulsion containing Carbohydrate fatty acid MonoSulphate derivative (CMS) induced the highest levels of cross-neutralizing antibodies compared to other oil-in-water emulsions or AS01B. Mechanistically, CMS induced antigen retention in the draining lymph node (dLN) and expression of cytokines, chemokines and type I interferon-stimulated genes at both injection site and dLN. Overall, CMS:RBD-NP is effective across multiple age groups and is an exemplar of a SARS-CoV-2 subunit vaccine tailored to the elderly.

## INTRODUCTION

SARS-CoV-2 emerged as a novel betacoronavirus at the end of 2019 and has rapidly spread throughout the world, leading to >150 million cases of Coronavirus Disease 2019 (COVID-19) globally including >32 million cases in the US (https://coronavirus.jhu.edu/). This pandemic has generated a global health crisis that will likely be ended only by widespread deployment of effective vaccines across all ages. Historically, the process of vaccine development spans several years. Basic research into prototype betacoronavirus pathogens coupled with advancements in structure-based antigen design, protein engineering and new manufacturing platforms such as mRNA and adenoviral virus vectors have enabled development of effective vaccines at an unprecedented speed (Gebre et al., 2021; Graham, 2020). Nevertheless, controlling the spread of the virus worldwide and especially in low- and middle-income countries will likely require global deployment of safe, effective, affordable, scalable, and practical vaccines that can also protect highly vulnerable populations including elderly individuals against COVID-19-related morbidity and mortality (Katz et al., 2021; Koff et al., 2021; Lancet Commission on and Therapeutics Task Force, 2021; Mejia et al., 2020). Protein subunit vaccines may meet at least some of these criteria and offer further advantages of not requiring ultra-cold storage and have a long track-record of safety. Indeed, SARS-CoV-2 subunit vaccines have already shown promising results in pre-clinical and clinical studies (Keech et al., 2020; Pollet et al., 2021; Richmond et al., 2021; Tian et al., 2021).

Most vaccines currently in use or in clinical development target the SARS-CoV-2 Spike glycoprotein due to the key role of its receptor-binding domain (RBD) in binding to the human receptor angiotensin-converting enzyme 2 (ACE2) and mediating cell entry (Lan et al., 2020; Walls et al., 2020b; Yan et al., 2020). The RBD protein would be an ideal candidate for a subunit vaccine since it is targeted by neutralizing antibodies (Abs) that exert a protective role against SARS-CoV-2 infection and it is readily produced at scale (Chen et al., 2021; Dalvie et al., 2021a; Dalvie et al., 2021b; Piccoli et al., 2020; Premkumar et al., 2020; Yang et al., 2020). However, RBD is poorly immunogenic and therefore its use as a candidate antigen poses limitations for vaccine development. To overcome this issue, two non-mutually exclusive approaches can be employed: 1) addition of adjuvant formulations, which are vaccine components that can enhance antigen immunogenicity by activating the innate immune system and/or modulating antigen pharmacokinetics (Irvine et al., 2020; Nanishi et al., 2020; O’Hagan et al., 2020; Pulendran et al., 2021) and 2) structure-based antigen design (Brune and Howarth, 2018; Graham et al., 2019; Irvine and Read, 2020; Kwong et al., 2020; Lopez-Sagaseta et al., 2016; Singh, 2021; Ward and Wilson, 2020). Regarding the latter, RBD antigens have been generated as dimers, trimers or displayed onto protein or synthetic nanoparticles with the goal of increasing antigen trafficking to the draining lymph node (dLN) and/or promoting clustering and activation of the B cell receptor (Arunachalam et al., 2021; Cohen et al., 2021; Dai et al., 2020; Dalvie et al., 2021b; Hauser et al., 2020; He et al., 2021; King et al., 2021; Ma et al., 2020; Sahin et al., 2020; Saunders et al., 2021; Tan et al., 2021; Walls et al., 2020a; Walsh et al., 2020). These approaches have successfully improved RBD immunogenicity and some of them are currently being evaluated in clinical trials (NCT04750343, NCT04742738). Assessing optimal combinations of candidate RBD antigens and adjuvants as well as their evaluation in pre-clinical models that take into account age-dependent vaccine immune responses and COVID-19 susceptibility will be key to down-selecting and prioritizing novel adjuvanted RBD antigen-based vaccines.

Here, we generated a nanoparticle in which multimeric RBD is displayed onto a protein scaffold composed of 60 subunits of the self-assembling bacterial protein lumazine synthase (Zhang et al., 2001). The RBD nanoparticle (RBD-NP) demonstrated higher immunogenicity compared to pre-fusion stabilized Spike trimer (Wrapp et al., 2020) (hereafter Spike) or monomeric RBD. By comparing multiple adjuvant formulations in combination with RBD-NP we found that a squalene-based oil-in-water (O/W) emulsion containing synthetic Carbohydrate fatty acid MonoSulphate derivative (CMS) (Hilgers et al., 2017) further enhanced anti-RBD serum Ab titers and SARS-CoV-2 cross-neutralizing titers in both young and aged mice. Mechanistically, CMS induced antigen retention in the dLN and expression of pro-inflammatory cytokines at the injection sites and type I interferon (IFN)-dependent IFN stimulated genes (ISGs) in the dLN. Overall, our study demonstrates the utility of a systematic and iterative approach to develop and optimize an adjuvanted RBD-NP effective across multiple age groups and provides further insights into the development of affordable SARS-CoV-2 subunit vaccines tailored to be active in the most vulnerable - i.e., the elderly.

## RESULTS

### *In vitro* characterization of RBD-NP reveals high density display multimeric RBD

High density display of antigens onto protein NPs increases their immunogenicity and has been employed in several vaccine candidates against viral infections to elicit robust serum antigen-specific Ab titers (Singh, 2021). In order to assemble SARS-CoV-2 RBD onto a protein NP scaffold we took advantage of the SpyTag/SpyCatcher conjugation system in which proteins fused with SpyTag and SpyCatcher spontaneously form stable isopeptide bonds (Brune et al., 2016). Briefly, we used the self-assembling lumazine synthase (LuS) from the hyperthermophile “*Aquifex aeolicus*” as protein NP scaffold (Zhang et al., 2001), and respectively expressed RBD and LuS with SpyCatcher (RBD-Catch) and SpyTag (LuS-Tag). SDS-PAGE analysis under reducing conditions of RBD-Catch, LuS-Tag, RBD-NP (generated by conjugating RBD-Catch and LuS-Tag) as well as native RBD and Spike proteins confirmed expected molecular weights (**Fig. 1A**). Transmission electron microscopy analysis of RBD-NP revealed a ruffled border, suggesting efficient conjugation and display of RBD onto the protein scaffold, and homogeneous size (**Fig. 1B**). This impression was further confirmed by dynamic light scattering analysis, showing an average size of ~30 nm (**Fig. 1C**). To confirm proper display of RBD onto NPs, we coated ELISA plates with RBD, Spike, RBD-NP, LuS-Tag, and assessed binding to recombinant human ACE2 (hACE2) and two anti-RBD monoclonal Abs (mAbs: clones H4 and CR3022). RBD-NP binding to hACE2, clones H4 and CR3022 was comparable to RBD and Spike while no binding to LuS-Tag was observed (**Fig. 1D, E**). RBD-NP binding profiles remained unaltered under multiple storage conditions, namely five freeze/thaw cycles or storage for 1 week at 4°C or room temperature (**Supplementary Fig. 1**). Interestingly, by assessing binding to mAb clones H4 and CR3022 under lower coating concentrations of RBD, Spike and RBD-NP we observed preferential binding to RBD-NP (**Fig. 1F**), suggesting that high density display of RBD onto NP increases Ab avidity. To explore another functional correlate of RBD-NP structure, we performed a competition assay in which Vero cells were incubated with SARS-CoV-2 in the absence or presence of multiple concentrations of RBD, Spike or RBD-NP. RBD-NP significantly reduced SARS-CoV-2 infection as assessed by IC50 and AUC (**Supplementary Fig. 2**), further supporting high-density display of RBD onto NP.

**Figure 1.**
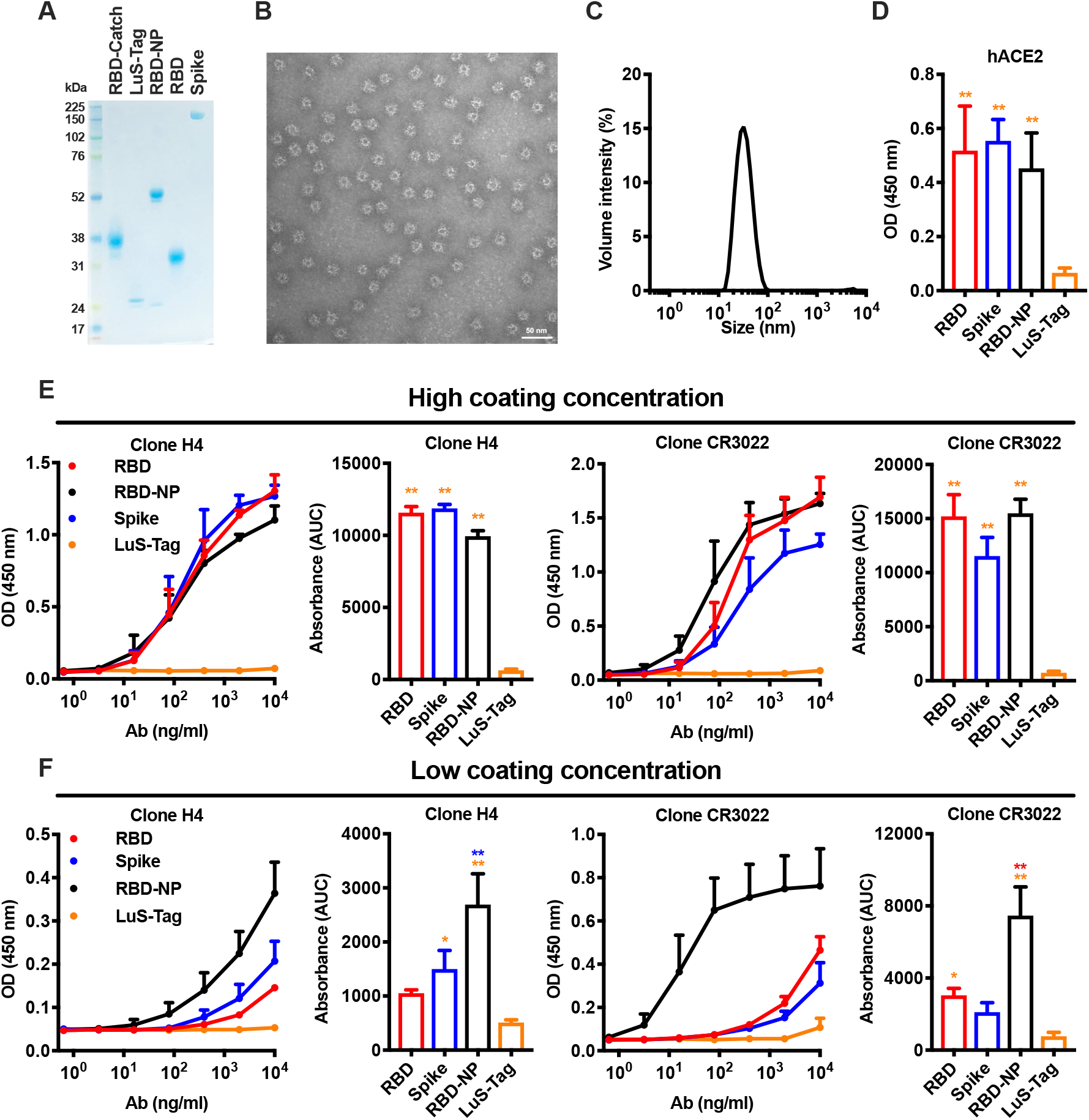
A lumazine synthase nanoparticle scaffold enables efficient RBD display. (**A**) SDS-PAGE analysis under reducing conditions of RBD expressing SpyCatcher (RBD-Catch), lumazine synthase expressing SpyTag (LuS-Tag), RBD nanoparticle (RBD-NP) as well as native RBD and Spike proteins. (**B**, **C**) Transmission electron microscopy (**B**) and dynamic light scattering (**C**) analyses of RBD-NP. (**D-F**) ELISA plates were coated with RBD, Spike, RBD-NP and LuS-Tag at 1 μg/ml (**D**), 5 μg/ml (**E**) or 0.5 μg/ml (**F**). Binding of recombinant human ACE2 (hACE2) or anti-RBD H4 and CR3022 antibody clones tested at multiple concentrations was expressed as optical density (OD) at 450 nm or area under the curve (AUC). N = 3-6 experiments. * and ** respectively indicate *p* ≤ 0.05 and 0.01. Statistical significance was determined by one-way ANOVA corrected for multiple comparisons. Comparisons among experimental groups are indicated by the color code.

### Immunization with RBD-NP elicits high serum anti-RBD antibody titers and SARS-CoV-2 neutralizing titers

To assess whether RBD-NP increases RBD immunogenicity,we immunized BALB/c mice with multiple doses of RBD, Spike or RBD-NP, alone or formulated with the MF59-like O/W emulsion *AddaVax* using a prime (Day 0) - boost (Day 14) schedule (**Fig. 2**). As expected, formulation with *AddaVax* enhanced anti-RBD Ab titers compared to immunizations with non-formulated antigens. In all experimental conditions, RBD-NP induced highest titers of anti-RBD IgG,IgG1 and IgG2a especially at the lowest tested dose (0.3 μg), thus showing a robust dose-sparing effect. Of note, anti-RBD Abs elicited by immunization with RBD-NP also recognized native RBD on Spike (**Supplementary Fig. 3**), which is key for SARS-CoV-2 neutralization. To confirm this point, we performed a surrogate of virus neutralization test (sVNT) that measure the degree of inhibition of RBD binding to hACE2 by immune sera, as well as a neutralization assay with live SARS-CoV-2 virus. In both assays, immunization with RBD-NP formulated with *AddaVax* induced higher levels of neutralization compared to immunization with Spike (**Fig. 3A, B**), while immunization with monomeric RBD failed to elicit significant levels of neutralization in the sVNT (**Fig. 3A**). Of note, immunization with RBD-NP formulated with *AddaVax* elicited high levels of anti-RBD neutralizing Abs in one additional inbred (C57BL/6) and one outbred (CD-1) mouse strains (**Supplementary Fig. 4**). Overall, these results show that multimeric RBD displayed on a NP significantly enhances its immunogenicity across multiple mouse strains, eliciting high levels of anti-RBD neutralizing Abs with significant dose-sparing effect.

**Figure 2.**
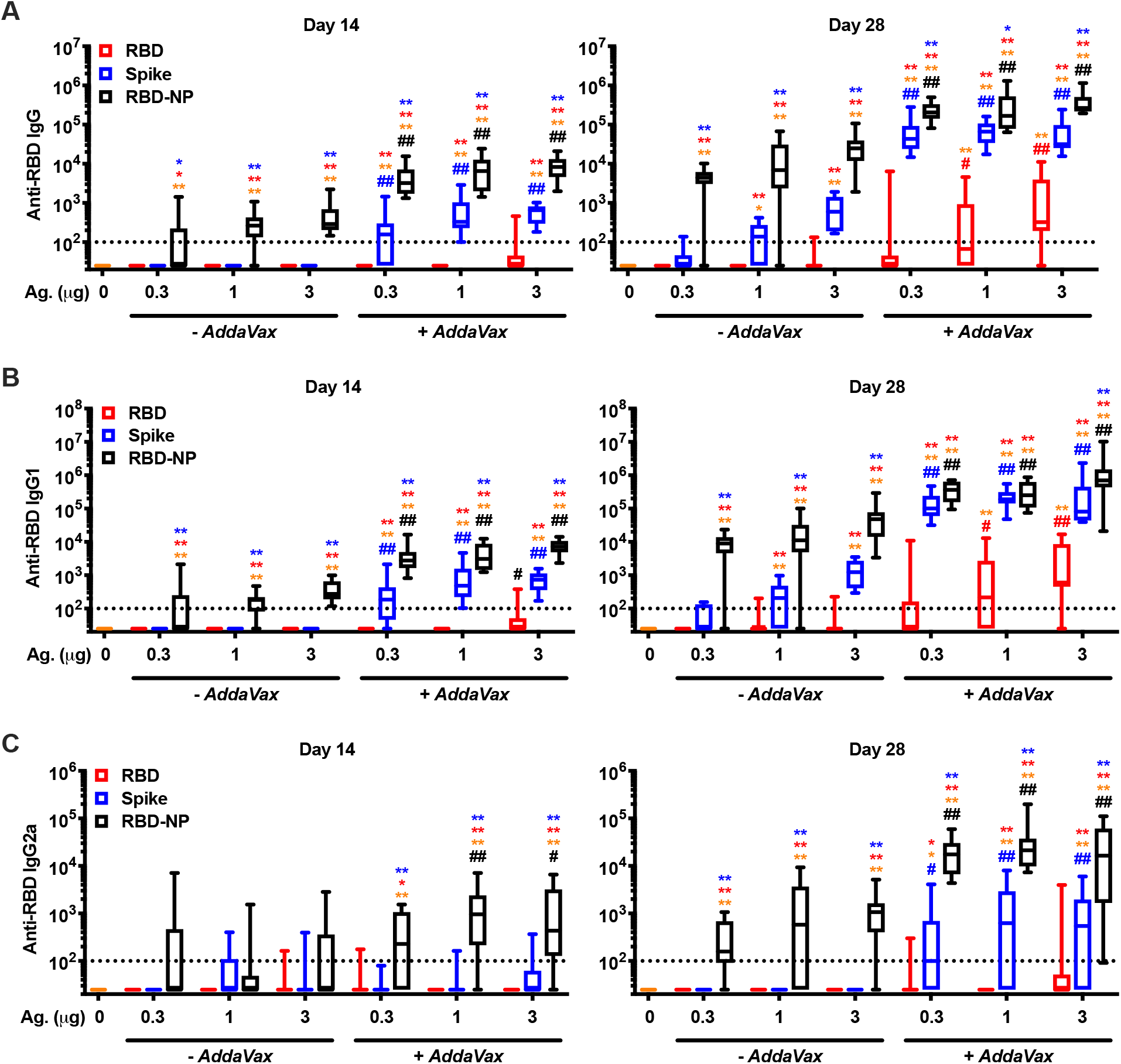
RBD nanoparticle demonstrates superior immunogenicity to Spike or monomeric RBD in mice. 3-month-old BALB/c mice were injected with PBS or immunized with the indicated doses of RBD, Spike or RBD nanoparticle (RBD-NP), alone or formulated with *AddaVax* on day 0 (prime) and 14 (boost). Anti-RBD IgG (**A**), IgG1 (**B**) and IgG2a (**C**) antibody titers were assessed in serum samples collected on days 14 (pre-boost) and 28. Dotted lines indicate lower limit of detection. N = 7-10 mice per group. * and ** respectively indicate *p* ≤ 0.05 and 0.01 for comparisons among RBD, Spike and RBD-NP in the same adjuvant formulation group (-*AddaVax* or +*AddaVax*). # and ## respectively indicate *p* ≤ 0.05 and 0.01 for comparisons of same antigen groups between the two adjuvant formulation groups (-*AddaVax* vs +*AddaVax*). Statistical significance was determined by two-way ANOVA corrected for multiple comparisons after Log-transformation of the raw data. Comparisons among experimental groups are indicated by the color code.

**Figure 3.**
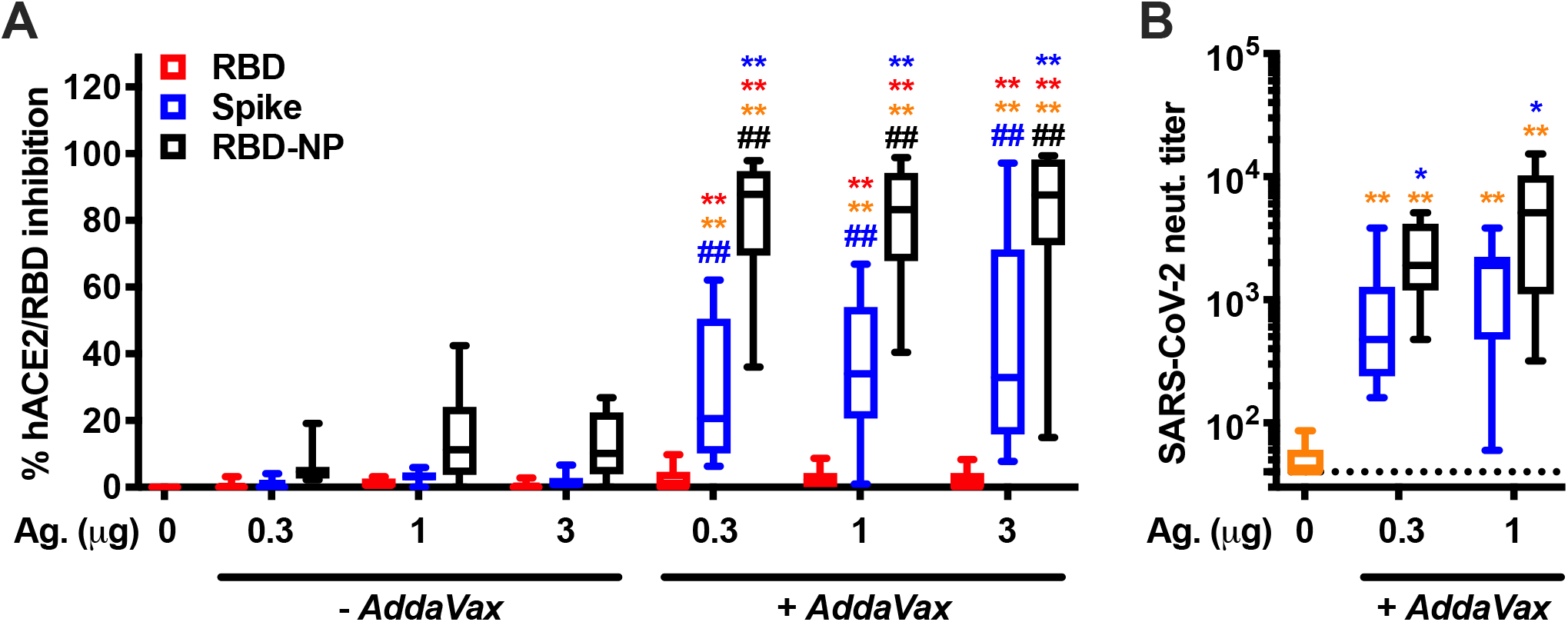
Immunization with RBD nanoparticle induces robust SARS-CoV-2 neutralizing titers at all doses tested. 3-month-old BALB/c mice were immunized as in **Figure 2**. Serum levels of anti-RBD neutralizing antibodies were assessed on day 28 by SARS-CoV-2 surrogate (**A**) and conventional (**B**) virus neutralization tests. The dotted line indicates lower limit of detection. N = 7-10 mice per group. * and ** respectively indicate *p* ≤ 0.05 and 0.01 for comparisons among RBD, Spike and RBD-NP in the same adjuvant formulation group (-*AddaVax* or +*AddaVax*). # and ## respectively indicate *p* ≤ 0.05 and 0.01 for comparisons of same antigen groups between the two adjuvant formulation groups (-*AddaVax* vs +*AddaVax*). Statistical significance was determined by two-way ANOVA corrected for multiple comparisons. Data shown in (**B**) were Log-transformed before the analysis. Comparisons among experimental groups are indicated by the color code.

### Novel CMS adjuvant significantly enhances RBD-NP immunogenicity in young and aged mice

Adjuvants play a key role in enhancing antigen immunogenicity (Irvine et al., 2020; Nanishi et al., 2020; O’Hagan et al., 2020; Pulendran et al., 2021). Although it is possible to generalize desirable properties for an adjuvant, a proper match between an adjuvant and a specific antigen has to be empirically evaluated. For example, comparisons of O/W- and aluminum hydroxide-based adjuvant formulations with RBD-NP demonstrated that the alpha-tocopherol-containing squalene-based oil-in-water emulsion AS03 is particularly effective at enhancing RBD-NP immunogenicity in non-human primates (Arunachalam et al., 2021). We therefore evaluated the immunogenicity of our RBD-NP with several O/W emulsions, namely *AddaVax*, the AS03-like adjuvant *AddaS03*, and a novel squalene-based O/W emulsion containing Carbohydrate fatty acid MonoSulphate derivative (CMS) (Hilgers et al., 2017). As a key benchmark, we also included AS01B (a liposome-based adjuvant containing monophosphoryl lipid A and saponin QS-21) as a clinical-grade benchmark adjuvant with potent immunostimulatory activity (Cunningham et al., 2016; Lal et al., 2015). As experimental model we chose to immunize both young (3-month-old) and aged (14-month-old) mice since we wanted to assess whether an optimized vaccine formulation could overcome impaired vaccine immunogenicity associated with immunosenescence in aged populations (Gustafson et al., 2020). All adjuvanted RBD-NP vaccine formulations induced robust titers of anti-RBD neutralizing Abs in young mice (**Fig. 4A-E**). Of note, CMS-adjuvanted RBD-NP vaccine elicited the highest levels of anti-RBD IgG Abs (**Fig. 4A**) by enhancing both anti-RBD IgG and IgG2a titers (**Fig. 4B, C**), resulting in potent inhibition of RBD binding to hACE2 (**Fig. 4D**) and SARS-CoV-2 neutralization (**Fig. 4E**). Immunization of aged mice resulted in overall lower anti-RBD Ab titers compared to young mice (**Supplementary Fig. 5**). Nevertheless, CMS again induced the highest anti-RBD antibody titers (**Fig. 4F-H**), inhibition of RBD binding to hACE2 (**Fig. 4I**) and SARS-CoV-2 neutralization (**Fig. 4J**) in aged mice.

**Figure 4.**
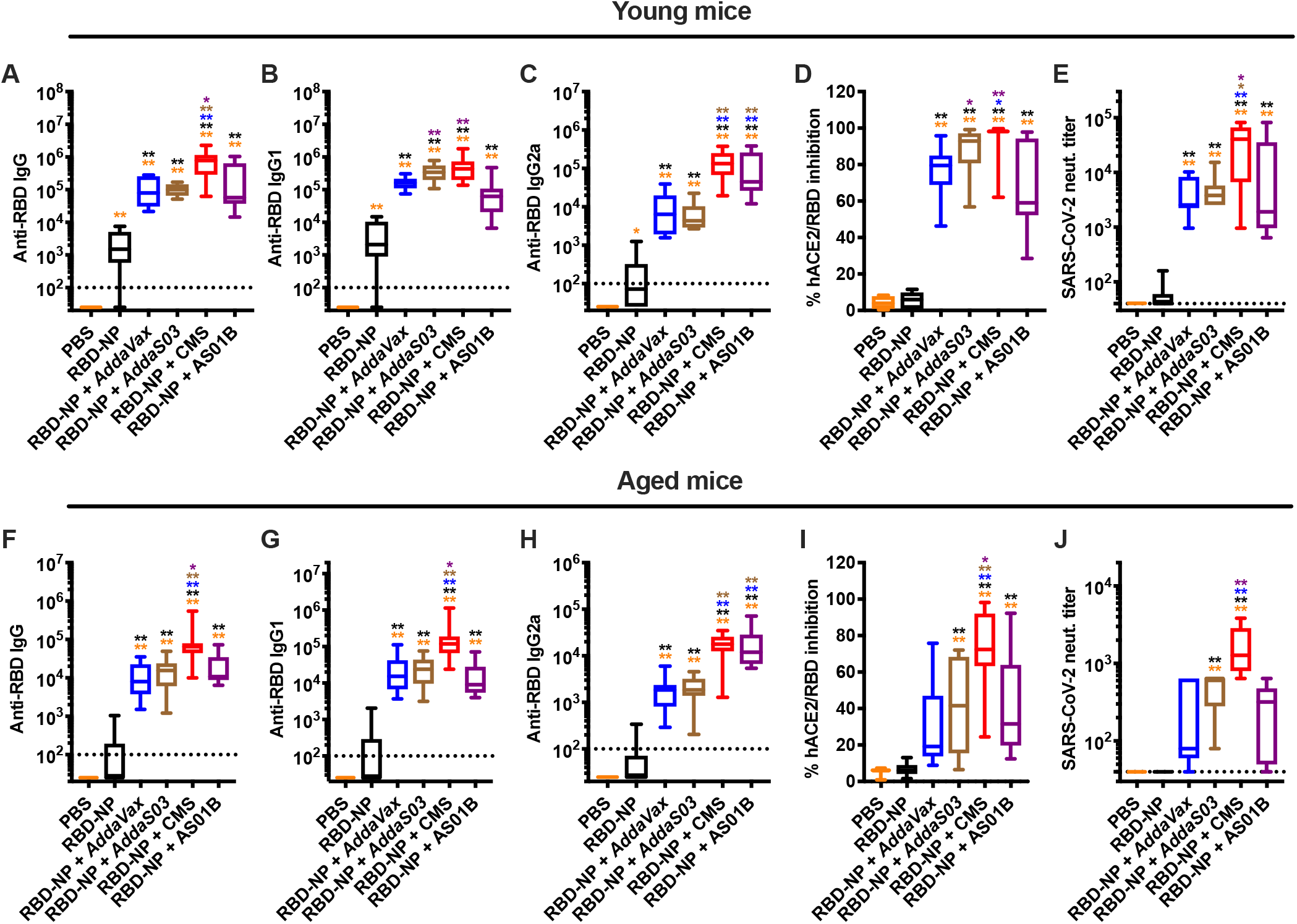
Formulation with CMS adjuvant enhances RBD nanoparticle immunogenicity in young and aged mice. Young (3-month-old,**A - E**) and aged (14-month-old,**F - J**) BALB/c mice were immunized as in **Figure 2** with PBS, RBD nanoparticle (RBD-NP) alone or formulated with *AddaVax, AddaS03*, CMS adjuvant or AS01B. Serum samples were collected on Day 28 to assess anti-RBD IgG (**A, F**), IgG1 (**B, G**), IgG2a (**C, H**) antibody titers as well as anti-RBD neutralizing activity by surrogate (**D, I**) and conventional (**E, J**) virus neutralization tests. Dotted lines indicate lower limit of detection. N = 10 (**A - I**) or 5 (**J**) mice per group. * and ** respectively indicate *p* ≤ 0.05 and 0.01. Statistical significance was determined by one-way ANOVA corrected for multiple comparisons. Data shown in (**A - C, E, F - I, J**) were Log-transformed before the analysis. Comparisons among experimental groups are indicated by the color code.

### Immunization with RBD-NP formulated with CMS adjuvant protects aged mice from SARS-CoV-2 challenge and elicits cross-neutralizing antibodies

Neutralizing Ab titers are an important correlate of protection against SARS-CoV-2 infection (Corbett et al., 2021; McMahan et al., 2021). As we observed high titers of neutralizing Abs in our immunization model, we decided to assess protection from SARS-CoV-2 infection by challenging immunized aged mice with 10^3^ PFU of the mouse adapted strain SARS-CoV-2 MA10 (Leist et al., 2020) and monitoring them up to four days post-infection (**Fig. 5**). Mice immunized with RBD-NP formulated CMS were fully protected from weight loss (**Fig. 5A**), and demonstrated low SARS-CoV-2 titers and *Il6, Ifit2* and *Rsad2* gene expression in the lungs (**Fig. 5B, C**).

**Figure 5.**
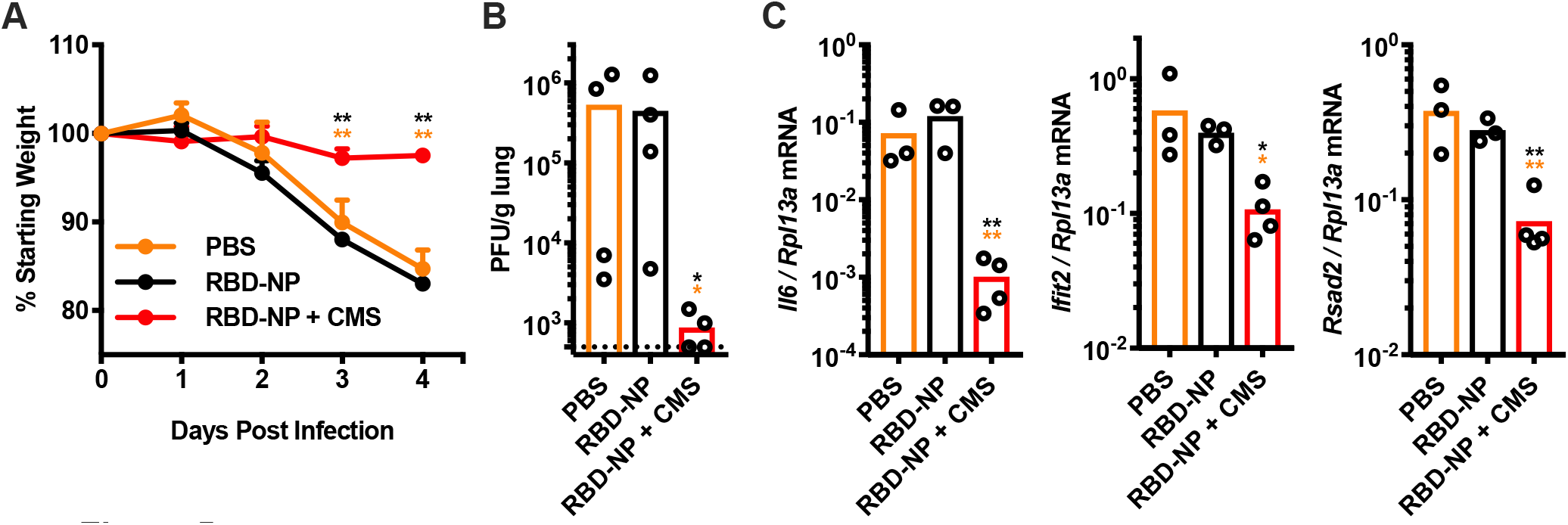
Immunization with CMS-adjuvanted RBD nanoparticle completely protects aged mice from SARS-CoV-2 challenge. (**A - C**) Aged (14-month-old) BALB/c mice were immunized as in **Figure 2** with PBS, RBD nanoparticle (RBD-NP) alone or formulated with CMS. On day 55 each mouse was infected with 10^3^ plaque-forming units (PFU) of mouse adapted SARS-CoV-2 and monitored up to 4 days for weight loss. (**A**) Daily weights of infected mice expressed as percentage of starting weight. (**B - C**) On Day 4, mice were sacrificed and lungs were collected to assess viral titers (**B**), expressed as PFU per gram of lung tissue) and gene expression profiles (**C**), shown as relative expression compared to *Rlp13a*.). N = 3-5 mice per group. Results are shown as mean + SEM (**A**) or as scatter dot plot and mean with each dot representing an individual sample (**B, C**). * and ** respectively indicate *p* ≤ 0.05 and 0.01. Statistical significance was determined by two-way ANOVA corrected for multiple comparisons (**A**) or one-way ANOVA corrected for multiple comparisons after Log-transformation of the raw data (**B, C**). Comparisons among experimental groups are indicated by the color code.

Since the beginning of the pandemic several SARS-CoV-2 variants such as B.1.17 and B.1.351 have emerged, the latter showing reduced neutralization by serum samples of convalescent or vaccinated subjects (Garcia-Beltran et al., 2021; Kuzmina et al., 2021; Shen et al., 2021). Therefore, we assessed neutralization of SARS-CoV-2 wild type (WT), B.1.17 and B.1.351 pseudoviruses by serum samples collected from young and aged mice immunized with RBD-NP formulated with *AddaVax, AddaS03*, CMS or AS01B (**Fig. 6**). As expected, we observed lower neutralizing titers against B.1.351 compared to WT. However, mice immunized with RBD-NP formulated with CMS showed the highest geometric mean titers against all SARS-CoV-2 variants, expecially in aged mice.

**Figure 6.**
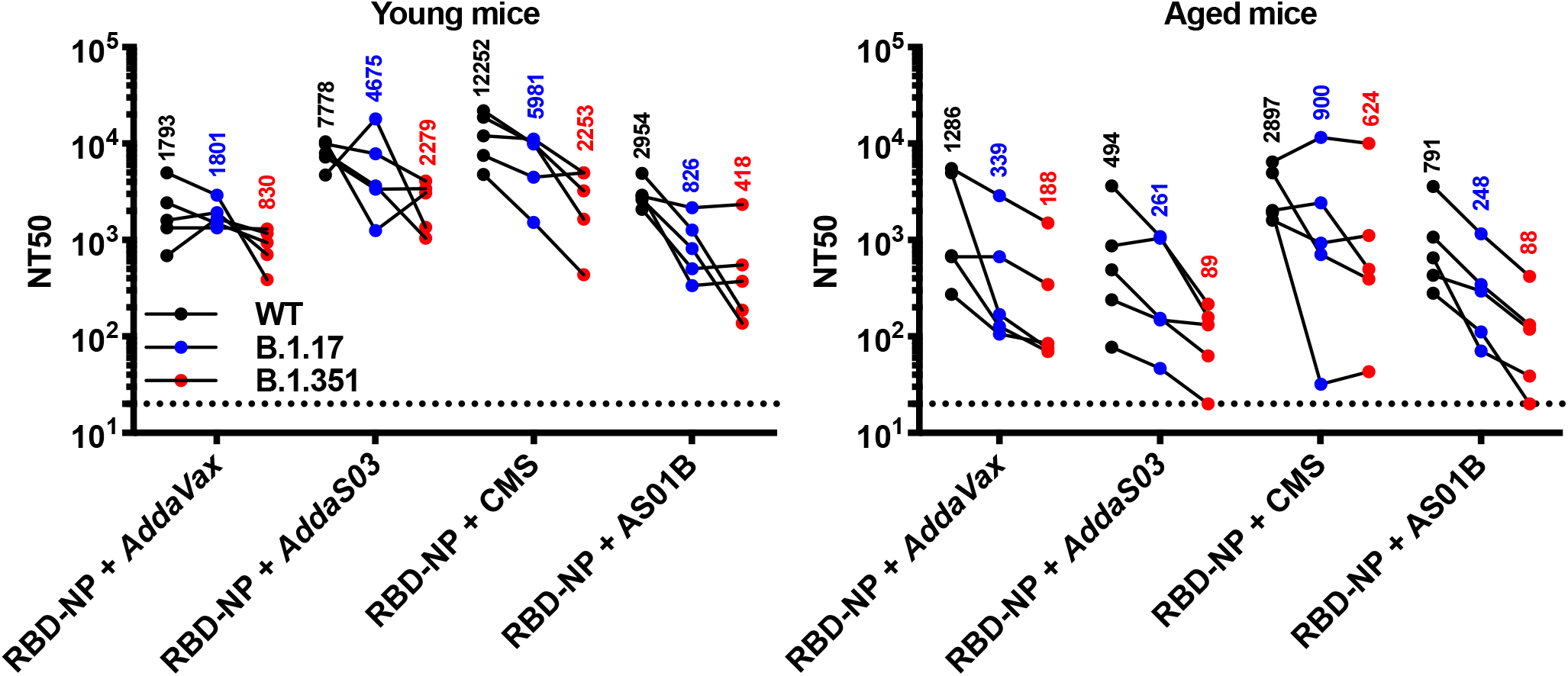
Immunization with CMS-adjuvanted RBD nanoparticle induces cross-neutralizing antibodies. Young (3-month-old) and aged (14-month-old) BALB/c mice were immunized as in **Figure 2** with RBD nanoparticle (RBD-NP) formulated with *AddaVax, AddaS03*, CMS adjuvant or AS01B. Serum samples were collected on Day 28 to assess neutralizing titers (NT50) agasint SARS-CoV-2 wild type (WT), B.1.17 or B.1.351 pseudoviruses. Dotted lines indicate lower limit of detection. N = 5 mice per group, with each dot representing an individual sample. Numbers indicate geometric mean titers for each experimental group.

### CMS adjuvant promote antigen retention in the draining lymph node

O/W emulsions are highly effective adjuvants and act through multiple mechanisms, including: 1) induction of a pro-inflammatory milieu at the injection site (Mosca et al., 2008) and/or 2) antigen targeting to and retention in the dLN (Cantisani et al., 2015). We therefore assessed whether the enhanced adjuvanticity of CMS could be explained by any of these two mechanisms. To this end, we injected mice with R-Phycoerythrin (R-PE) as a model protein antigen with intrinsic fluorescence, alone or formulated with CMS or *AddaVax* that we used as benchmark adjuvant. Twenty four hours post-injection, both *AddaVax* and CMS promoted significant and comparable antigen retention in the dLN (Supplementary Fig. 6). Interestingly, both adjuvants induced high gene expression of pro-inflammatory cytokines (*Csf2, Il6, Cxcl1*) and ISGs (*Cxcl9, Ifit2, Rsad2*) at the injection site, with *AddaVax* further enhancing the expression of the latter (Fig. 7A). However, only CMS enhanced type I IFN-dependent ISG expression in the dLN (Fig. 7B).

**Figure 7.**
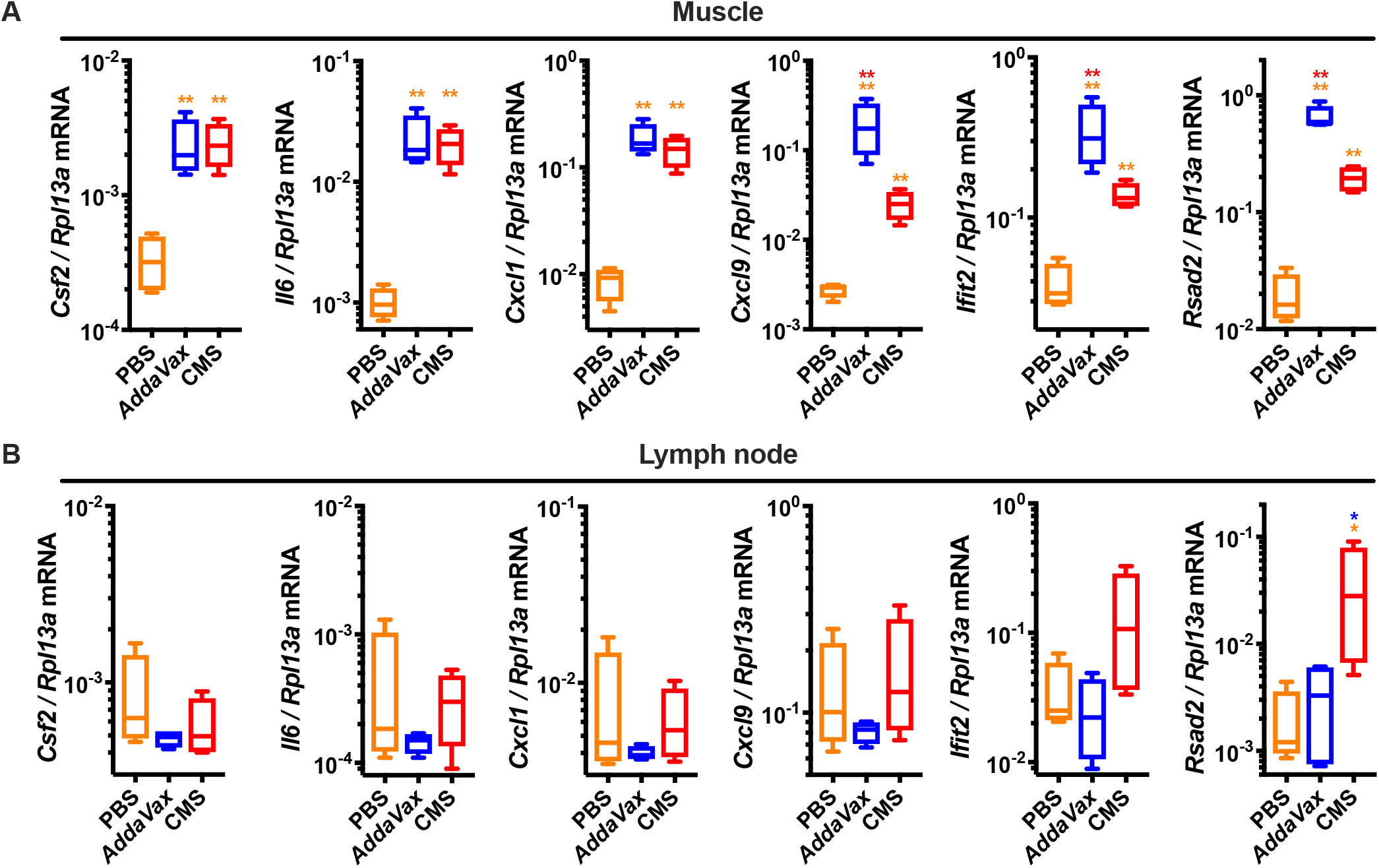
CMS adjuvant induces distinct gene expression profiles from *Addavax*,including enhanced type I IFN-dependent ISG expression in the draining lymph node. (**A, B**) Young (3-month-old) BALB/c mice were injected with PBS, *AddaVax* and CMS adjuvant. 24 hours later muscle tissue at the injection sites (**A**) and dLNs (**B**) were collected to assess gene expression profiles by qPCR. Results are reported as relative expression compared to *Rlp13a*. N = 4 mice per group. * and ** respectively indicate *p* ≤ 0.05 and 0.01. Statistical significance was determined by one-way ANOVA corrected for multiple comparisons after Log-transformation of the raw data. Comparisons among experimental groups are indicated by the color code.

## DISCUSSION

Despite rapid development and deployment of effective SARS-CoV-2 vaccines based on mRNA or viral vector technologies (Gebre et al., 2021; Graham, 2020), there remains an urgent global need for affordable, scalable, and practical coronavirus vaccines (Katz et al., 2021; Koff et al., 2021; Lancet Commission on and Therapeutics Task Force, 2021; Mejia et al., 2020). In this context, adjuvanted protein subunit vaccines are likely to play an important role for increasing worldwide vaccine coverage, based in part on their generally robust history of safety and efficacy in special populations such as the infant and the elderly. Adjuvanted subunit vaccines consisting of Spike or RBD proteins, expressed in soluble or nanoparticle forms, are currently in different stages of pre-clinical and clinical development (Arunachalam et al., 2021; Cohen et al., 2021; Dai et al., 2020; Dalvie et al., 2021b; Hauser et al., 2020; He et al., 2021; Keech et al., 2020; King et al., 2021; Ma et al., 2020; Pollet et al., 2021; Richmond et al., 2021; Saunders et al., 2021; Tan et al., 2021; Tian et al., 2021; Walls et al., 2020a). On-going efforts are required to define optimal combinations of antigens and adjuvants to enhance immunogenicity across the lifespan. Here, we assessed the immunogenicity of RBD-NP formulated with four different adjuvants, namely the MF59-like *AddaVax*, AS03-like *AddaS03*, AS01B, and CMS. Consistent with prior studies, we found that the first three elicited high levels of anti-RBD neutralized Abs (Arunachalam et al., 2021; King et al., 2021; Walls et al., 2020a). However, CMS was the most effective at enhancing RBD-NP immunogenicity in both young and aged mice and protected the latter from live SARS-CoV-2 challenge. To provide a mechanistic basis for this phenomenon, we found that CMS promotes antigen retention in the dLN and induced a gene expression profiles at both the injection site and the dLN that are distinct from the ones elicited by *AddaVax*, including induction of type I IFN-dependent ISGs in the dLN that may promote immunogenicity (De Giovanni et al., 2020). Overall, our study provides novel mechanistic and translational information on O/W adjuvants and might inform the development of adjuvanted RBD-NP vaccines effective across multiple age groups.

In this study, we compared the immunogenicity of monomeric RBD, Spike and RBD-NP using the same adjuvant formulation, namely *AddaVax*. As expected, monomeric RBD was poorly immunogenic. RBD-NP was more immunogenic than Spike across all tested dosed. Our results are supported by the concept that antigen display onto NP scaffold enhances immunogenicity and activate B cells to eventually produce antigen-specific Abs (Brune and Howarth, 2018; Graham et al., 2019; Irvine and Read, 2020; Kwong et al., 2020; Lopez-Sagaseta et al., 2016; Singh, 2021; Ward and Wilson, 2020). However, a recent study has reported that immunizations of non-human primates with RBD-NP or Spike Hexapro formulated with AS03 elicited comparable SARS-CoV-2 neutralizing titers (Arunachalam et al., 2021). Whether the differences between these findings and our study are due to use of different reagents (distinct RBD-NP designs, Spike vs Spike Hexapro, AddaVax vs AS03), doses and/or animal models (mice vs non-human primates) has not yet been determined. Nevertheless, both studies as well as additional recent publications support the use of RBD-NP as an effective SARS-CoV-2 vaccine antigen (Arunachalam et al., 2021; Cohen et al., 2021; Dalvie et al., 2021b; He et al., 2021; King et al., 2021; Ma et al., 2020; Saunders et al., 2021; Tan et al., 2021; Walls et al., 2020a).

To further optimize RBD-NP immunogenicity we compared four adjuvant formulations, namely *AddaVax, AddaS03*, AS01B and CMS. The first three (or similar adjuvant formulations) increase RBD-NP immunogenicity (Arunachalam et al., 2021; King et al., 2021; Walls et al., 2020a), with an AS03-adjuvanted RBD-NP being currently evaluated in a clinical trial (NCT04750343). CMS is a potent adjuvant with a low reactogenicity profile (Hilgers et al., 2017) but until now had never been tested with an RBD-NP. The CMS adjuvant technology is a third generation of the so-called carbohydrate fatty acid sulphate esters-based adjuvants such as CoVaccine HT (Blom and Hilgers, 2004). Compared to CoVaccine HT, CMS has an improved safety profile but similar abilities to induce high antibody titers. Strikingly, CMS induced the highest levels of anti-RBD IgG Abs in young and aged mice by enhancing both anti-RBD IgG1 and IgG2a. This translated into high SARS-CoV-2 neutralizing titers that protected aged mice from SARS-CoV-2 challenge. Serum samples of mice immunized with RBD-NP formulated with CMS also demonstrated the highest neutralization titers against B.1.17 and B.1.351 pseudoviruses which represent SARS-CoV-2 variants of concern. Therefore, CMS is a promising adjuvant formulation for an RBD-NP-based SARS-CoV-2 vaccine effective across multiple age groups.

To define a mechanistic basis for CMS adjuvanticity, we hypothesized that CMS could induce significant local cytokine and chemokine production at the injection site and/or antigen retention in the dLN. We focused on these mechanisms as they have been described previously for O/W emulsions (Cantisani et al., 2015; Mosca et al., 2008). By comparing CMS and *AddaVax* adjuvants, we found that both induced high antigen retention in the dLN and expression of pro-inflammatory genes at the injection site. Cytokine and chemokine production at the injection site can promote innate immune cell recruitment, activation and subsequent antigen presentation (Mosca et al., 2008). However, only CMS enhanced type I IFN-dependent ISG expression in the dLN which can promote differentiation of CD4^+^ T follicular helper cells and therefore antigen-specific antibody response (De Giovanni et al., 2020). Further work will be required to define the precise mechanism of action of CMS, as well as how antigen localization and germinal center dynamics in the dLN are modulated by *AddaVax* and CMS.

The emergence of SARS-CoV-2 variants that can escape neutralizing Abs elicited by infection or vaccination has raised concerns regarding possible reduction in vaccine efficacy (Garcia-Beltran et al., 2021; Kuzmina et al., 2021; Shen et al., 2021). While our results show significant neutralization of B.1.17 and B.1.351 pseudoviruses by serum samples of mice immunized with RBD-NP and CMS, it is likely that novel vaccines specifically targeting SARS-CoV-2 variants of concern may need to be rapidly developed (Wu et al., 2021). In addition, the risk posed by emerging zoonotic coronaviruses calls for the development of pan-coronavirus vaccines that can protect against circulating variants as well as strains currently circulating only in non-human animals but that can generate future human outbreaks and pandemics. The SpyTag/SpyCatcher conjugation platform employed in our study has recently been used to generate mosaic NP displaying RBD proteins derived from up to 8 strains (Cohen et al., 2021). Whether different adjuvant formulations (e.g. *AddaVax*, CMS) can modulate the breadth of the Ab response elicited by mosaic RBD-NP is an important area for future research.

Overall, we have described the process of selection and optimization of an adjuvanted RBD-NP vaccine formulation capable of inducing cross-neutralizing Abs and protective immunity across multiple age groups, even at very low doses, and provided a mechanistic basis for its enhanced immunogenicity. Our work may inform further pre-clinical and clinical development of precision adjuvanted RBD-NP-based SARS-CoV-2 and pan-coronavirus vaccines tailored for robust immunogencity and protection in the vulnerable elderly.

## ACKNOWLEDGEMENTS

We thank the members of the BCH *Precision Vaccine Program* for helpful discussions. We thank Drs. Kevin Churchwell, Gary Fleisher, David Williams, and Mr. August Cervini for their support of the *Precision Vaccines Program*. We thank Dr. Barney S. Graham (NIH Vaccine Research Center) for generously providing the plasmid for pre-fusion stabilized SARS-CoV-2 Spike trimer. We thank Dr. Ralph Baric for providing the SARS-CoV-2/MA10 virus. D.J.D. would like to thank Ms. Siobhan McHugh, Ms. Geneva Boyer, Mrs. Lucy Conetta and the staff of Lucy’s Daycare, the staff the YMCA of Greater Boston, Bridging Independent Living Together (BILT), Inc., and the Boston Public Schools for childcare and educational support during the COVID-19 pandemic. The current study was supported in part by US National Institutes of Health (NIH)/National Institutes of Allergy and Infectious Diseases (NIAID) awards, including Adjuvant Discovery (HHSN272201400052C and 75N93019C00044) and Development (HHSN272201800047C) Program Contracts to O.L. as well as a Massachusetts Consortium on Pathogenesis Readiness (MassCPR) grant to A.O., L.R.B., and O.L. D.J.D.’s laboratory is supported by NIH grant (1R21AI137932-01A1), Adjuvant Discovery Program contract (75N93019C00044). REH is supported by the U.S. Army’s Long Term Health and Education Training Program and MBF is partially supported by BARDA# ASPR-20-01495, DARPA# ASPR-20-01495, NIH R01 AI148166, and NIH HHSN272201400007C. A.G.S.’s laboratory is supported by NIH R01 AI146779 (A.G.S.), NIGMS T32 GM007753 (B.M.H. and T.M.C.), T32 AI007245 (J.F.), and a Massachusetts Consortium on Pathogenesis Readiness (MassCPR) grant to A.G.S. I.Z. is supported by NIH grants 1R01AI121066 and 1R01DK115217, and holds an Investigators in the Pathogenesis of Infectious Disease Award from the Burroughs Wellcome Fund. The *Precision Vaccines Program* is supported in part by the BCH Department of Pediatrics and the Chief Scientific Office and acknowledges philanthropic support via the BCH Trust. E.N. is supported by the Daiichi Sankyo Foundation of Life Science and Uehara Memorial Foundation and is a joint Society for Pediatric Research and Japanese Pediatric Society Scholar.

## AUTHOR CONTRIBUTIONS

FB and EN conceived, designed, performed, analyzed the experiments and wrote the paper; TRO and YS performed *in vitro* and/or *in vivo* experiments and their analysis; JC, JD-A and AO contributed to data analysis; KS, AZX, HSS and SDP expressed and purified SARS-CoV-2 RBD, Spike and RBD-NP; TMC, JF, BMH and AGS provided SARS-CoV-2 RBD plasmid, expressed and purified SARS-CoV-2 RBD; MEM, REH, CD, SMW, HLH, RMJ, and MBF performed and analyzed SARS-CoV-2 neutralization experiments and mouse challenge study; AC, JY and DHB performed pseudovirus neutralization assays; PPP and LH provided CMS adjuvant and contributed to the design of experiments; RKE and IZ contributed to the design of experiments; OL and DJD contributed funding, assisted with study design, supervised the study and edited the manuscript.

## METHODS

### Protein expression and purification

Full length SARS-CoV-2 spike glycoprotein (M1-Q1208, GenBank MN90894) and RBD constructs (amino acid residues R319-K529, GenBank MN975262.1), both with an HRV3C protease cleavage site, a TwinStrepTag and an 8XHisTag at C-terminus, were obtained from Barney S. Graham (NIH Vaccine Research Center) and Aaron Schmidt (Ragon Institute), respectively. To generate RBD-Catch and LuS-Tag constructs in a mammalian expression vector, sARS-CoV-2 RBD and SpyCatcher (Brune et al., 2016) were fused by a GGSGGS linker for RBD-Catch, and N-terminal Spy-tag was added to lumazine synthase from *Aquifex aeolicus* bearing D71N mutation for LuS-Tag. Both contructs contain a signal peptide (MKHLWFFLLLVAAPRWVLS) at N-terminus and HRV3C protease site, followed by a TwinStrepTag at C-terminus. These mammalian expression vectors were used to transfect Expi293F suspension cells (Thermo Fisher) using polyethylenimine (Polysciences). Cells were allowed to grow in 37°C, 8% CO_2_ for additional 5 days before harvesting for purification. Protein was purified in a PBS buffer (pH 7.4) from filtered supernatants by using either StrepTactin resin (IBA) or Cobalt-TALON resin (Takara). Affinity tags were cleaved off from eluted protein samples by HRV 3C protease, and tag removed proteins were further purified by size-exclusion chromatography using a Superose 6 10/300 column (Cytiva) for full length Spike and a Superdex 200 10/300 Increase 10/300 GL column (Cytiva) for RBD, RBD-Catch, and LuS-Tag in a PBS buffer (pH 7.4).

### RBD-Catch and LuS-Tag conjugations

To saturate Lus-Tag surface with RBD, a 1:1.2 molar ratio of LuS-Tag and RBD-Catch components were mixed at 40 μM of LuS-Tag in a PBS buffer (pH 7.4) and incubated at room temperature for approximately 1 hour. Reaction mixture was applied to a Superdex200 Increase 10/300 GL column (Cytiva) in a PBS buffer (pH 7.4) to purify RBD-nanoparticles from unconjugated RBD-Catch. The conjugated RBD-nanoparticle product was confirmed by SDS-PAGE and analyzed by negative-stain EM.

### SDS-PAGE analysis

Proteins samples (250 μg/ml) in NuPAGE LDS Sample Buffer (Invitrogen) were heated to 95°C for 5 min and 10 μl (2.5 μg) were loaded to a NuPAGE 10% Bis-Tris gel (Invitrogen). The gel was run in NuPAGE MOPS Buffer (Invitrogen) at 60 V for 45 min and then 110 V for 105 min. The gel was then rinsed with DI water and fixed for 15 min in 50 mL of 40% ethanol, 10% acetic acid fixing solution. The gel was rinsed with DI water and incubated with QC Colloidal Coomassie Blue (Bio-Rad) on a rotating shaker for 1 hour at RT. The gel was rinsed twice with DI water and incubated on a rotating shaker for 75 min, changing the water every 15 min, and then imaged.

### Negative staining electron microscopy

Purified RBD-nanoparticle samples were diluted to 0.01-0.05 mg/mL with a PBS (pH 7.4) buffer. A 4-μl drop of the diluted sample was applied to a freshly glow-discharged carbon-coated copper grid (400-mesh, EMS) for approximately one minute. The drop was removed using blotting paper, and the grid was washed three times with 5-μl drops of the same buffer. Adsorbed proteins were negatively stained by soaking in 4-μl drops of 2% uranyl acetate for approximately 10 s and removing drop with filter paper. Micrographs were collected using JEM-1400 Plus electron microscope (JEOL, USA) operated at 80 kV, resulting in ~0.15nm/pixel at 80,000x magnification.

### Dynamic light scattering

Purified protein samples (250 μg/ml) were loaded into disposable microcuvette and measured at 25 °C using a Zetasizer Ultra instrument (Malvern Panalytical) equipped with a 633-nm laser with 3 scans of 60 sec each. Each sample was measured in triplicate, and the intensity of the size distribution was plotted in GraphPad Prism 9 (GraphPad Software).

### Enzyme-Linked Immunosorbent Assay (ELISA)

ELISA was employed to examine the binding ability of the purified proteins to hACE2 and RBD-specific monoclonal Abs (mAbs). Briefly, RBD monomer, Spike trimer, RBD-Catch, LuS-Tag, and RBD-NPs were respectively diluted to concentrations of 0.5 and 5 μg/ml for mAb binding and 1 μg/ml for hACE2 binding, and 50 μl/well were added to coat 96-well high-binding flat-bottom plate (Corning) overnight at 4 °C. Plates were washed with 0.05% Tween 20 PBS (PBS-T) and blocked with 1% BSA PBS for 1 h at room temperature (RT). The plates were then incubated with sequentially 1:5 serially diluted hACE2-Fc (invivogen) and two anti-SARS-CoV-2 RBD mAbs (clones H4 [InvivoGen] and CR3022 [Abcam]) starting at 10 μg/ml in blocking buffer. After 2 h of incubation, the plates were washed with PBS-T three times, and incubated with HRP-conjugated detection Abs (mouse anti-human IgG1 Fc-HRP, Southern Biotec). Plates were washed five times and developed with tetramethylbenzidine (OptEIA Substrate Solution, BD Biosciences) for 5 min, then stopped with 2 N H_2_SO_4_. Optical densities (ODs) were read at 450 nm with SpectraMax iD3 microplate reader (Molecular Devices).

### RBD-NP stability analysis

RBD-NP samples (1 mg/ml) were subjected to one to five cycles of freeze–thaw cycles by storing in a −80 °C freezer for at least 1 day, followed by incubation at RT for 30 min. For the storage temperature study, RBD-NP samples (1 mg/ml) were incubated at 4 °C or RT for 5-7 days. The RBD-NP samples were then analyzed by ELISA.

### Live SARS-CoV-2 *in vitro* competition assay

The day prior to infection, 5e3 VeroE6 cells were plated per well in DMEM (Quality Biological) supplemented with 10% v/v fetal bovine serum (Gibco), 1% v/v Penicillin-Streptomycin (Gemini-Bio), and 1% v/v L-Glutamine (Gibco). 1 mg/ml stock concentrations of SARS-CoV-2 Spike, SARS-CoV-2 RBD, and SARS-CoV-2 RBD-NP were diluted to 50 μg/ml in 400 μl complete VeroE6 media in a 96-well dilution block in duplicate and then serially diluted down the plate 1:3 to produce an 8-point dilution curve (125 μl into 250 μl media). Media was removed from the VeroE6 cells and 90 μl of each dilution was then transferred to the cells and left to incubate at 37°C and 5% CO_2_ for 2 hours. After incubation, each well was infected with a 0.1 M.O.I. of SARS-CoV-2 △ORF7a::GFP (provided by Dr. Ralph Baric (UNC)) diluted in 10 μl media. A parallel plate was left uninfected to monitor cytotoxicity. After 48 hours, the infected plates were fixed in 4% paraformaldehyde for 1 hour, Hoechst-stained, and read on a plate reader (Nexcelom Biosciences, Lawrence, MA). The percentage of GFP+ cells in each well was counted and compared to an untreated, infected control to give an inhibitory concentration 50 (IC_50_) for each protein. The parallel cytotoxicity plate was analyzed with Cell Titer Glo (Promega, Madison, WI) and read on a BioTek Synergy HTX plate reader (BioTek Instruments, Inc., Winooski, VT). Cell viability was compared to an untreated control.

### Animals

Female, 3 month old BALB/c and C57BL/6J mice were purchased from Jackson Laboratory (Bar Harbor, ME), and CD-1 mice were purchased from Charles River Laboratories (Wilmington, MA). Female, 12-13 months old BALB/c mice purchased from Taconic Biosciences (Germantown, NY) were used for aged mice experiments. Mice were housed under specific pathogen-free conditions at Boston Children’s Hospital, and all procedures were approved under the Institutional Animal Care and Use Committee (IACUC) and operated under the supervision of the Department of Animal Resources at Children’s Hospital (ARCH) (Protocol number 19-02-3897R). At the University of Maryland School of Medicine, mice were housed in a biosafety level 3 (BSL3) facility for all SARS-CoV-2 infections with all procedures approved under the IACUC (Protocol number #1120004).

### Mouse immunization

All formulations for immunization were prepared under sterile conditions. Mice were injected with antigens (RBD monomer, Spike trimer, and RBD-NPs), with or without adjuvants. Mock treatment mice received phosphate-buffered saline (PBS) alone. Injections (50 μl) were administered intramuscularly in the caudal thigh on Days 0 and 14. The adjuvants and their doses used were: *AddaVax* (25 μl), *AddaS03* (25 μl) (InvivoGen), AS01B (40 μl) (obtained from the Shingrix vaccine, GSK Biologicals SA, Belgium), and CMS adjuvant (25 μl corresponding with 1 mg of CMS) (LiteVax, The Netherlands).

### Mouse serum antibody ELISA

RBD- and Spike-specific Ab titers were quantified in serum samples by ELISA by modifitation of a previously described protocol (Borriello et al., 2017). Briefly, high-binding flat-bottom 96-well plates (Corning) were coated with 50 ng/well RBD or 25 ng/well Spike and incubated overnight at 4 °C. Plates were washed with PBS-T (PBS + 0.05% Tween 20) and blocked with 1% BSA PBS for 1 h at RT. Serum samples were serially diluted 4-fold from 1:100 up to 1:1.05^8^ and then incubated for 2 hours at RT. Plates were washed three times and incubated for 1 hour at RT with HRP-conjugated anti-mouse IgG, IgG1, IgG2a, or IgG2c (Southern Biotech). Plates were washed five times and developed with tetramethylbenzidine (1-Step Ultra TMB-ELISA Substrate Solution, ThermoFisher, for RBD-ELISA,and BD OptEIA Substrate Solution, BD Biosciences, for Spike ELISA) for 5 min, then stopped with 2 N H_2_SO_4_. Optical densities (ODs) were read at 450 nm with SpectraMax iD3 microplate reader (Molecular Devices). End-point titers were calculated as the dilution that emitted an optical density exceeding a 3× background. An arbitrary value of 25 was assigned to samples with OD values below the limit of detection for which it was not possible to interpolate the titer.

### Surrogate of virus neutralization test (sVNT)

We performed sVNT to measure the degree of hACE2/RBD inhibition by immune sera, with modification of a previously published protocol (Tan et al., 2020). Briefly, high-binding flat-bottom 96-well plates (Corning, NY) were coated with 100 ng/well recombinant human ACE2 (hACE2) (Sigma-Aldrich) in PBS, incubated overnight at 4°C, washed three times with PBS-T, and blocked with 1% BSA PBS for 1 hour at RT. Each serum sample was diluted 1:160, pre-incubated with 3 ng of RBD-Fc in 1% BSA PBS for 1 hour at RT, and then transferred to the hACE2-coated plate. RBD-Fc without pre-incubation with serum samples was added as a positive control, and 1% BSA PBS without serum pre-incubation was added as a negative control. Plates were then washed three times and incubated with HRP-conjugated anti-human IgG Fc (Southern Biotech) for 1 hour at RT. Plates were washed five times and developed with tetramethylbenzidine (BD OptEIA Substrate Solution, BD Biosciences) for 5 min, then stopped with 2 N H_2_SO_4_. The optical density was read at 450 nm with SpectraMax iD3 microplate reader (Molecular Devices). Percentage inhibition of RBD binding to hACE2 was calculated with the following formula: Inhibition (%) = [1 – (Sample OD value – Negative Control OD value)/(Positive Control OD value – Negative Control OD value)] × 100.

### Live SARS-CoV-2 virus neutralization test

All serum samples were heat-inactivated at 56°C for 30 min to deactivate complement and allowed to equilibrate to RT prior to processing for neutralization titer. Samples were diluted in duplicate to an initial dilution of 1:20 followed by 1:2 serial dilutions (vaccinated samples), resulting in a 12-dilution series with each well containing 60 μl. All dilutions employed DMEM (Quality Biological), supplemented with 10% (v/v) fetal bovine serum (heat-inactivated, Gibco), 1% (v/v) penicillin/streptomycin (Gemini Bio-products) and 1% (v/v) L-glutamine (2 mM final concentration, Gibco). Dilution plates were then transported into the BSL-3 laboratory and 60 μl of diluted SARS-CoV-2 (WA-1, courtesy of Dr. Natalie Thornburg/CDC) inoculum was added to each well to result in a multiplicity of infection (MOI) of 0.01 upon transfer to titering plates. A non-treated, virus-only control and mock infection control were included on every plate. The sample/virus mixture was then incubated at 37°C (5.0% CO_2_) for 1 hour before transferring 100 μl to 96-well titer plates with 5e3 VeroE6 cells. Titer plates were incubated at 37°C (5.0% CO_2_) for 72 hours, followed by cytopathic effect (CPE) determination for each well in the plate. The first sample dilution to show CPE was reported as the minimum sample dilution required to neutralize >99% of the concentration of SARS-CoV-2 tested (NT99).

### Pseudovirusneutralization test

The SARS-CoV-2 pseudoviruses expressing a luciferase reporter gene were generated in an approach similar to as described previously (Yu et al., 2021; Yu et al., 2020). Briefly, the packaging plasmid psPAX2 (AIDS Resource and Reagent Program), luciferase reporter plasmid pLenti-CMV Puro-Luc (Addgene), and spike protein expressing pcDNA3.1-SARS CoV-2 SΔCT of variants were co-transfected into HEK293T cells by lipofectamine 2000 (ThermoFisher). Pseudoviruses of SARS-CoV-2 variants were generated by using WA1/2020 strain (Wuhan/WIV04/2019, GISAID accession ID: EPI_ISL_402124), B.1.1.7 variant (GISAID accession ID: EPI_ISL_601443), or B.1.351 variant (GISAID accession ID: EPI_ISL_712096). The supernatants containing the pseudotype viruses were collected 48 h post-transfection, which were purified by centrifugation and filtration with 0.45 μm filter. To determine the neutralization activity of the plasma or serum samples from participants, HEK293T-hACE2 cells were seeded in 96-well tissue culture plates at a density of 1.75 × 10^4^ cells/well overnight. Three-fold serial dilutions of heat inactivated serum or plasma samples were prepared and mixed with 50 μL of pseudovirus. The mixture was incubated at 37°C for 1 h before adding to HEK293T-hACE2 cells. 48 h after infection, cells were lysed in Steady-Glo Luciferase Assay (Promega) according to the manufacturer’s instructions. SARS-CoV-2 neutralization titers were defined as the sample dilution at which a 50% reduction in relative light unit (RLU) was observed relative to the average of the virus control wells.

### SARS-CoV-2 mouse challenge study

Mice were anesthetized by intraperitoneal injection of 50μL of a xylazine and ketamine mix (0.38 mg/mouse and 1.3 mg/mouse, respectively) diluted in PBS. Mice were then inoculated intranasally with 1 × 10^3^ PFU of mouse-adapted SARS-CoV-2 (MA10, courtesy of Dr. Ralph Baric (UNC)) in 50 μl divided between nares (Leist et al., 2020). Challenged mice were weighed on the day of infection and daily for up to 4 days post-infection. At 4-days post-infection, mice were sacrificed, and lungs were collected to assess virus load by plaque assay and gene expression profiles. SARS-CoV-2 lung titers were quantified by homogenizing harvested lungs in PBS (Quality Biological Inc.) using 1.0 mm glass beads (Sigma Aldrich) and a Beadruptor (Omni International Inc.). Homogenates were added to Vero E6 cells and SARS-CoV-2 virus titers determined by counting plaque-forming units (pfu) using a 6-point dilution curve. RNA was isolated from lung homogenates using Direct-zol RNA miniprep kit (Zymo Research), according to the manufacturer’s protocol. RNA concentration and purity (260/280 and 260/230 ratios) were measured by NanoDrop (ThermoFisher Scientific).

### Analysis of inflammatory responses at injection site and dLN

Young (3-month-old) BALB/c mice were injected with PBS, *AddaVax* or CMS adjuvant, and their local muscle tissue and dLN were harvested for subsequent analysis 24 hours later. For dLN analysis, adjuvants were injected in caudal thigh and inguinal LNs were collected. For muscle tissue analysis, adjuvants were injected in the gastrocnemius muscle, and whole gastrocnemius was collected. Samples were stored in RNA*later* (Invtrogen) for 24 hours at 4°C and then homogenized in TRI Reagent (Zymo Research) with a beadbeater. Samples were then centrifuged, and the clear supernatant was transferred to a new tube for subsequent RNA isolation. RNA was isolated from TRI Reagent samples using phenol-chloroform extraction or column-based extraction systems (Direct-zol RNA Miniprep, Zymo Research) according to the manufacturer’s protocol. RNA concentration and purity (260/280 and 260/230 ratios) were measured by NanoDrop (ThermoFisher Scientific).

### Gene expression analysis by qPCR

Purified RNA was analyzed for gene expression by qPCR on a CFX384 real time cycler (Bio-rad) using pre-designed KiCqStart SYBR Green Primers (MilliporeSigma) specific for *Csf2* (RM1_Csf2 and FM1_Csf2), *Cxcl9* (RM1_Cxcl9 and FM1_Cxcl9), *Ifit2* (RM1_Ifit2 and FM1_Ifit2), *Rsad2* (RM1_Rsad2 and FM1_Rsad2), *Il6* (RM1_Il6 and FM1_Il6), *Cxcl1* (RM1_Cxcl1 and FM1_Cxcl1), *Rpl13a* (RM1_Rpl13a and FM1_Rpl13a).

### Measurement of antigen retention within the dLN

We assessed antigen retention effect of indicated adjuvants, with modification of a previously published protocol (Cantisani et al., 2015). Briefly, young (3-month-old) BALB/c mice were injected IM with vaccine formulation (50 μL) applying R-phycoerythrin (R-PE) as a model antigen (6 μg). 24 hours later, the dLN was collected and homogenized in water with a beadbeater. Fluorescence values were measured with SpectraMax i3x microplate reader (Molecular Devices) and expressed as arbitrary units after background (deionized water) subtraction.

### Statistical analysis

Statistical analyses were performed using Prism v9.0.2 (GraphPad Software). Some datasets were analyzed after Log-transformation as indicated in the figure legends. Statistical differences between groups in datasets with one categorical variable were evaluated by two sample t test (2 groups) or one-way ANOVA (more than 2 groups) corrected for multiple comparisons. Statistical differences between groups in datasets with two categorical variables were evaluated by two-way ANOVA corrected for multiple comparisons. *p* values ≤ 0.05 were considered significant.

## SUPPLEMENTAL INFORMATION

**Supplementary Figure 1. RBD nanoparticle is stable under multiple storage conditions.** ELISA plates were coated with RBD nanoparticles that underwent 1 (F/T ×1) or 5 (F/T ×5) freeze/thaw cycles, or stored for 1 week at 4°C (4°C - 1wk) or room temperature (RT - 1wk). Binding of recombinant human ACE2 (hACE2) or anti-RBD H4 and CR3022 antibody clones was expressed as optical density (OD) at 450 nm or area under the curve (AUC). N = 4 experiments. Statistical significance was determined by one-way ANOVA corrected for multiple comparisons.

**Supplementary Figure 2. RBD nanoparticle competes with SARS-CoV-2 *in vitro*.** (**A, B**) Vero cells were infected with SARS-CoV-2 in the presence or absence of RBD, Spike and RBD nanoparticle (RBD-NP) tested at multiple concentrations. Results are expressed as percentage of non-infected cells. For each protein a non-linear curve was fitted and used to calculate IC50 (**A**) and area under the curve (**B**). N = 2 experiments.

**Supplementary Figure 3. Anti-RBD antibodies elicited by RBD nanoparticle immunization recognize native RBD on Spike.** Anti-Spike IgG, IgG1 and IgG2a antibody titers were assessed in serum samples collected on Day 28 as indicated in **Figure 2**. Dotted lines indicate lower limit of detection. N = 10 mice per group. ** indicates *p* ≤ 0.01. Statistical significance was determined by one-way ANOVA corrected for multiple comparisons after Log-transformation of the raw data. Comparisons among experimental groups are indicated by the color code.

**Supplementary Figure 4. RBD nanoparticle is immunogenic in multiple mouse strains.** 3-month-old C57BL/6 (**A**) and CD-1 (**B**) mice were injected with PBS or immunized with 0.3 μg RBD nanoparticle (RBD-NP), alone or formulated with *AddaVax* on Day 0 (prime) and Day 14 (boost). Anti-RBD IgG, IgG1 and IgG2a antibody titers were assessed in serum samples collected on Day 28. Dotted lines indicate lower limit of detection. N = 5 mcie per group. * and ** respectively indicate *p* ≤ 0.05 and 0.01. Statistical significance was determined by one-way ANOVA corrected for multiple comparisons after Log-transformation of the raw data. Comparisons among experimental groups are indicated by the color code.

**Supplementary Figure 5. Aged mice demonstrate reduced anti-RBD antibody response upon immunization.** Comparisons of anti-RBD IgG, IgG1 and IgG2a antibody titers between immunized young and aged mice as reported in **Figure 4**. N = 10 mice per group * and ** respectively indicate *p* ≤ 0.05 and 0.01. Statistical significance was determined by two-way ANOVA corrected for multiple comparisons after Log-transformation of the raw data. Comparisons among experimental groups are indicated by the color code.

**Supplementary Figure 6. *AddaVax* and CMS adjuvant promote antigen retention in the draining lymph node.** 3-month-old BALB/c mice were injected intramuscularly with PBS, R-PE, R-PE formulated with *AddaVax* or CMS adjuvant. 24 hours later draining lymph node were collected, homogenized in water and fluorescence was measured in cleared supernatants. Results are expressed as arbitrary units (A.U.) of fluorescence. N = 12 mice per group. ** indicates *p* ≤ 0.01. Statistical significance was determined by one-way ANOVA corrected for multiple comparisons after Log-transformation of the raw data. Comparisons among experimental groups are indicated by the color code.

