## Supplementary figures and images for "An adjuvanted SARS-CoV-2 RBD nanoparticle elicits neutralizing antibodies and fully protective immunity in aged mice"

### Supplementary Figure 1

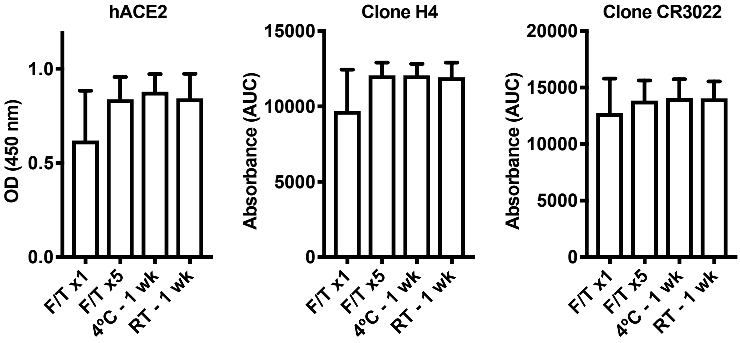


##### Supplementary Figure 1

### Supplementary Figure 2

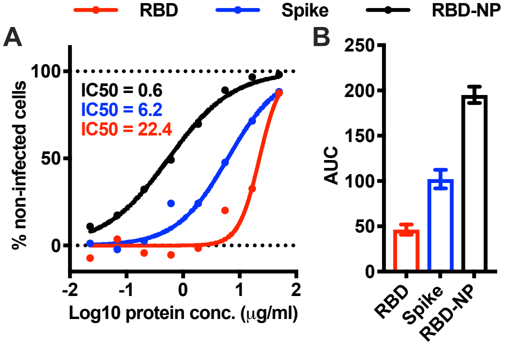


#### Supplementary Figure 2

### Supplementary Figure 3

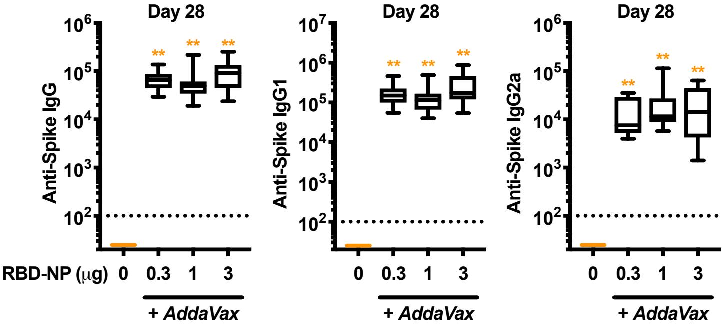


##### Supplementary Figure 3

### Supplementary Figure 4

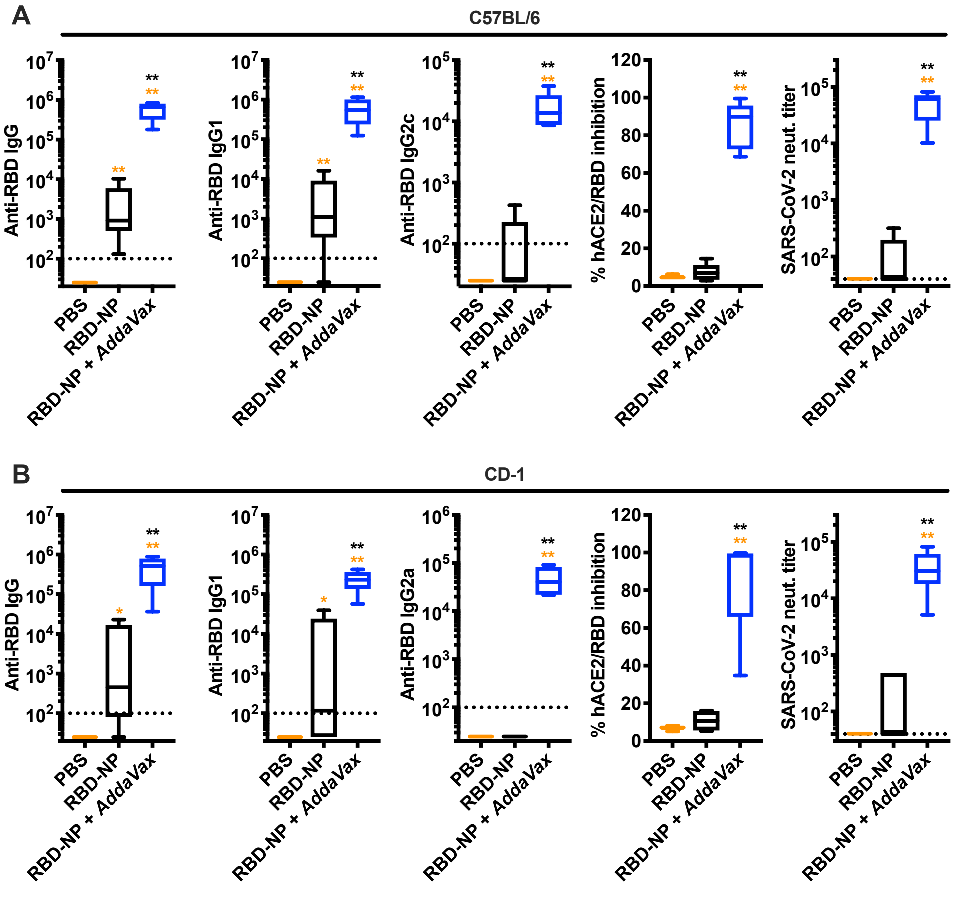


**Supplementary Figure 4**

### Supplementary Figure 5

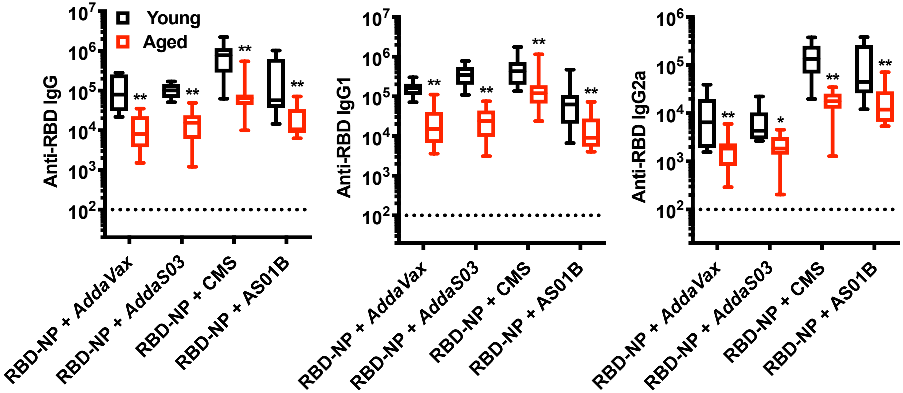


**Supplementary Figure 5**

### Supplementary Figure 6

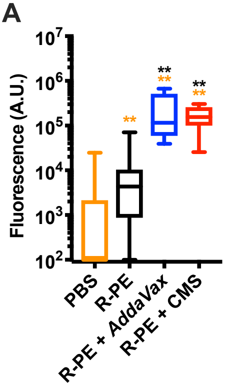


#### Supplementary Figure 6
